# Exploring Xylem Anatomical Adaptations Associated with Crassulacean Acid Metabolism and Hydraulic Capacitance in *Clusia* Leaves: Lessons for CAM Bioengineering

**DOI:** 10.1101/2022.05.27.493620

**Authors:** Alistair Leverett, Kate Ferguson, Klaus Winter, Anne M. Borland

**Affiliations:** School of Natural and Environmental Sciences, Newcastle University, Newcastle Upon Tyne, NE1 7RU, UK; Smithsonian Tropical Research Institute, PO Box 0843-03092, Balboa, Ancón, Republic of Panama

**Keywords:** Crassulacean Acid Metabolism, Hydraulic Capacitance, Xylem, Vein Density, Vessel, *Clusia*, Leaf Anatomy

## Abstract

**Background and Aims:** Many succulent species are characterised by the presence of crassulacean acid metabolism (CAM) and/or elevated bulk hydraulic capacitance (C_FT_). Both CAM and elevated C_FT_ substantially reduce the rate at which water moves through transpiring leaves. However, little is known about how these physiological adaptations are coordinated with leaf vascular architecture and xylem anatomy.

**Methods:** The genus *Clusia* contains species spanning the entire C_3_-CAM continuum, and also is known to have > 5-fold interspecific variation in C_FT_. We used this highly diverse genus to explore how interspecific variation in vein density and xylem vessel dimensions are coordinated with CAM and C_FT_.

**Key Results:** We found that constitutive CAM phenotypes were associated with lower vein length per leaf area (VLA) and vein termini density (VTD), compared to C_3_ or facultative CAM species. However, when vein densities were standardised by leaf depth, this value was higher in CAM than C_3_ species, which is likely an adaptation to overcome apoplastic hydraulic resistance in deep chlorenchyma tissue. In contrast, C_FT_ did not correlate with any xylem anatomical trait measured, suggesting CAM has a greater impact on leaf hydraulic physiology than C_FT_.

**Conclusions:** Our findings strongly suggest that CAM photosynthesis is coordinated with leaf vein densities. The link between CAM and vascular anatomy will be important to consider when attempting to bioengineer CAM into C_3_ crops.

## Introduction

The field of leaf hydraulics has grown in the last 15 years, following the discovery that leaves act as a hydraulic bottleneck in trees, accounting for more than 30 % of the hydraulic resistance in the soil-plant-atmosphere continuum (Sack et al., 2003; Sack & Holbrook, 2006; Sack & Tyree, 2005; Scoffoni et al., 2011). More than 40 trillion tonnes of water traverse plants each year, and with the growing threat of global warming it is essential that we understand the relationships between physiological traits and leaf xylem anatomy (Chahine, 1992; Sack & Tyree, 2005; Sheffield & Wood, 2008; Choat et al., 2018; Jiao et al., 2021). Crassulacean acid metabolism (CAM) is a photosynthetic adaptation that minimises water loss via reorganised stomatal dynamics, so that stomata open at night and close during the middle of the day, (Borland *et al*., 2014, 2015; Yang *et al*., 2015). By decreasing transpiration, CAM prevents leaf water potentials (Ψ_L_) from falling low enough to cause mechanical damage to vascular and mesophyll tissues (Haag-Kerwer *et al*., 1996; Winter *et al*., 2005; Leverett *et al*., 2022). Elevated bulk hydraulic capacitance (C_FT_: see equation 1 in Materials and Methods) is another strategy for coping with water limitation which allows leaves to maintain a stable Ψ_L_ when tissues dehydrate (Smith *et al*., 1987; Ogburn and Edwards, 2010, 2012; Luo *et al*., 2021). Consequently, leaves with high C_FT_ rely less on the influx of water to prevent transpiration-induced drops in Ψ_L_ (Leverett *et al*., 2022). Hence, like CAM, elevated C_FT_ minimises the rate at which water flows through transpiring leaves. Despite the fact that both CAM and elevated C_FT_ reduce the flux of water through transpiring leaves, little is known about how these traits are coordinated with vascular architecture and xylem anatomical characteristics (Males, 2017b; Borland *et al*., 2018). Understanding how CAM and leaf vascular traits are coordinated is important for informing attempts to bioengineer the CAM pathway into C_3_ crops. Work is already underway to select C_3_ host plants with succulent mesophyll anatomy, in order to ensure the efficient function of bioengineered CAM pathways (Lim *et al*., 2018, 2020). However, little consideration has been given to the xylem traits that will be needed to optimise the hydraulics in CAM leaves. By identifying ways in which CAM and xylem anatomy are co-adapted, our findings will aid bioengineering efforts to select/design appropriate host plants with optimal vascular architecture.

To address the relationship between leaf vascular traits CAM and C_FT_, we focused on the genus *Clusia*. These tropical trees, epiphytes and hemiepiphytes exhibit a wide variety of photosynthetic phenotypes, including obligate C_3_ and constitutive CAM species as well as C_3_-CAM intermediates, where CAM accounts for only a fraction of total carbon assimilation (Barrera-Zambrano *et al*., 2014; Borland *et al*., 2018; Leverett *et al*., 2021; Luján *et al*., 2021; Pachon *et al*., 2022). In addition, a > 5-fold variation in C_FT_ exists across *Clusia* (Leverett *et al*., 2022). CAM and C_FT_ appear to be independent traits in this genus, as the former is associated with deep chlorenchyma tissue, whereas the latter is the consequence of investment in adaxial hypodermal hydrenchyma tissue (Leverett *et al*., 2022). As CAM and C_FT_ are independent adaptations in *Clusia* leaves, this genus is ideal for exploring which trait is more influential on vascular architecture and xylem anatomy.

Xylem conduits within leaves consist of narrow tracheids and wider vessels which, together, conduct water into the distal portions of the lamina. The vasculature of eudicots is typically arranged in a single plane in a leaf, such that larger veins furcate into smaller, finer veins that radiate into the leaf lamina (Esau, 1965; Nelson and Dengler, 1997). The midrib and the two orders of veins that branch from this are known as major veins, and are characterised by thick, structurally reinforced xylem tissue. The finer, branched veins that furcate off the third order veins (i.e. fourth order and above) are described as minor veins; these typically radiate into the lamina tissue until they eventually end at vein termini (Sack and Scoffoni, 2013). Vein length per area of leaf (VLA), determines a species’ ability to conduct water across the leaf (Boyce *et al*., 2009; Scoffoni *et al*., 2011; Sack and Scoffoni, 2013). A higher VLA has two complementary effects: it facilitates the flow of water inside the xylem, whilst also minimising the distance water needs to travel outside of the xylem (Scoffoni *et al*., 2017). Consequently, species with higher VLA can more easily conduct water across their leaves (Scoffoni *et al*., 2016, 2017). As CAM and elevated C_FT_ both decrease the rate of water moving into and across leaves, we hypothesised that species with these adaptations would have lower VLA.

Whilst it is useful to consider vasculature in a two-dimensional plane, leaves are in fact 3 dimensional organs, and the 3-D density of veins is determined by both VLA and the depth of the mesophyll tissue in which they are found (Zwieniecki and Boyce, 2014; Males, 2017a). Noblin *et al*., (2008) used both model and real leaves to show that optimal vascular architecture occurs when intervein distance (IVD) roughly equals the vein to lower epidermal distance (IVD ≈ VED). An IVD:VED ratio > 1 means that insufficient veins were available to efficiently replace the water that was lost from stomata. Conversely, an IVD:VED ratio < 1 would mean that superfluous veins are present that don’t increase the efficiency of mesophyll hydraulic conductance. Across diverse angiosperm species, IVD and VED were found to be approximately equal, reinforcing the argument that this arrangement is optimal in leaves (Zwieniecki and Boyce, 2014). However, some species with thicker leaves overinvest in veins, meaning they have IVD:VED ratios < 1 (de Boer *et al*., 2016; Males, 2017a). Two opposing hypotheses have been proposed to explain this. One suggestion is that greater leaf thickness causes higher hydraulic resistance in the mesophyll apoplast, meaning that leaves need overinvestment in vein placement to maintain n efficient hydraulic conductance across the whole leaf (de Boer *et al*., 2016). A contrasting suggestion is that high C_FT_ in thick leaves could require greater vascular conductance in order to quickly refill water reserves following rainfall (Males, 2017a). *Clusia* is an ideal model to test these two alternative hypotheses, as C_FT_ is independent of CAM and leaf thickness in this genus. Therefore, it is possible to investigate if high IVD:VED ratios are determined more by CAM/leaf thickness or by C_FT_/hydrenchyma depth.

Finally, to complete the characterisation of vascular traits associated with CAM, we investigated xylem anatomy inside the veins, by measuring the dimensions of the water-conducting vessels. Water travels across the xylem conduits under tension, due to the pulling force from transpirational water loss (Sack and Holbrook, 2006; Brodribb *et al*., 2016). Under tension, water exists at a metastable state and can spontaneously change phase from liquid to gas, which breaks the contiguity of the transpiration stream and prevents hydraulic conductance (Wagner *et al*., 2022). Wider xylem conduits are more vulnerable to the formation of emboli, which can ‘seed’ the formation of gaseous bubbles and thereby curtail hydraulic conductance (Lens *et al*., 2013; Scoffoni *et al*., 2016). Consequently, a trade-off exists, whereby wider vessels allow more water to flow into leaves, but these same wide vessels are also more vulnerable to drought-induced embolism. However, both CAM and elevated C_FT_ slow the speed at which Ψ_L_ falls, thereby preventing emboli from forming. Therefore, we hypothesised that by buffering Ψ_L_ and protecting leaves from forming emboli, CAM and elevated C_FT_ would enable species to evolve wider vessels to more efficiently conduct water. Since first order major veins are known to be the predominant site at which emboli originate, (Brodribb *et al*., 2016; Hochberg *et al*., 2016; Klepsch *et al*., 2018) we measured vessel dimensions in the petiole and central midrib across the selected *Clusia* species.

Based on the aforementioned considerations we conducted interspecific comparisons across the genus *Clusia* to test three hypotheses:

1. Are CAM and/or elevated C_FT_ associated with lower VLA?
2. Are CAM and/or elevated C_FT_ associated with IVD:VED ratios < 1?
3. Are CAM and/or elevated C_FT_ associated with wider vessels in the midrib and petiole?

By testing these hypotheses, we were able to generate a fundamental understanding of the ways in which vascular architecture and xylem anatomy are coordinated with CAM and C_FT_ in leaves.

## Materials and Methods

### Species Studied

This comparative study investigated a well-characterised and phylogenetically diverse collection of *Clusia* species (Gustafsson *et al*., 2007; Barrera-Zambrano *et al*., 2014 - and references therein; Leverett *et al*., 2021). Species studied reflected a diversity of photosynthetic phenotypes: obligate C_3_ species – *Clusia multiflora, C. tocuchensis* and *C. grandiflora*; the intermediate/facultative CAM species – *C. lanceolata, C. aripoensis, C. pratensis* and *C. minor*; and constitutive CAM species – *C. rosea, C. fluminensis, C. alata* and *C. hilariana*.

### Plant Growth Conditions

Plants were grown in a glasshouse in Cockle Park farm, as a part of Newcastle University’s *Clusia* collection. The 3-6-year-old plants (approx. 60-100 cm tall) were grown in 3:1, (v/v) compost-sand mixture (John Innes No. 2, Sinclair Horticulture Ltd, Lincoln, UK), in 10 L pots. The glasshouse has fitted photosynthetic LED lights (Attis 5 LED plant growth light, PhytoLux) allowing plants to receive a minimum 12-hour light day. The glasshouse temperatures were 25 °C during the day and 23 °C at night.

### Photosynthetic Gas Exchange

Gas exchange data used in these comparative analyses was extracted from Barrera-Zambrano *et al*. (2014) along with data on *C. pratensis* and *C. fluminensis* from Leverett *et al*. (2021). Net CO_2_ uptake was recorded over 24 hours, using a BINOS infrared gas analyser (Walz). The proportion of total diel assimilation occurring during the night in well-watered plants (CAM_ww_) and after 9 days of drought (CAM_d_) was used as a quantitative estimate of CAM.

### Leaf Water Relations

Estimates of leaf bulk hydraulic capacitance (C_FT_) were taken from Leverett *et al*. (2022). Pressure-volume curve data were used to calculate C_FT_ with the equation

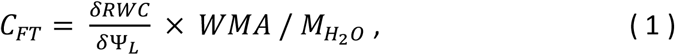

where RWC is relative water content, Ψ_L_ is leaf water potential, WMA is the mass of water per leaf area of fully hydrated leaves, and M_H2O_ is the molar mass of water.

### Vein Length per Leaf Area (VLA) and Intervein Distance (IVD)

Some succulent species are known to have veins organised in multiple planes within a leaf, a phenomenon known as 3D vasculature (Balsamo and Uribe, 1988; Cutler, 2004; Ogburn and Edwards, 2013; Heyduk *et al*., 2016; Males, 2017a; Fradera-Soler *et al*., 2021; Jolly *et al*., 2021). Before measuring VLA, leaves were hand-sectioned, to visually check the arrangement of veins. 3D vasculature was not observed in any species of *Clusia* used in this study (Fig. S1). Vein length per leaf area (VLA) was measured according to the protocol described in Scoffoni *et al*., (2011), with modifications for working with *Clusia* leaves. From each plant, the fourth leaf from the apex was sampled and leaf area measured using a flatbed scanner (hp Scanjet 5530 Photosmart Scanner, Hp, UK). Primary VLA (i.e. VLA of the midrib) was measured by dividing the length of the midrib by the leaf area. Due to the thick, waxy leaves of *Clusia* it was not possible to clear whole leaves to measure VLA, so instead, for each leaf, a rectangle (2 × 3 cm, or 1 by 2 cm for smaller leaves of *C. lanceolata* and *C. minor)* was cut halfway along the proximal-distal axis of the leaf blade. This rectangle did not include the leaf margin or the midrib. Both abaxial and adaxial surfaces were gently rubbed with a nail file to remove some wax and make fine perforations. Leaf tissue was soaked for 45 minutes in 3:1 95 % ethanol:acetic acid (Fisher Chemical) to further remove wax. Pigments were then cleared by soaking tissue in 5 % NaOH (w/v) (BDH Chemicals Ltd) for 45 minutes. Leaf material was transferred to 50 % (v/v) bleach in aqueous solution for 15 minutes to remove blackened phenolics. Leaf material was washed in water 4 times, each for 15 minutes, and then dehydrated by transferal to solutions containing increasing concentrations of ethanol (dehydration series was 30 %, 50 %, 70 % and 100 % ethanol, each lasting 20 minutes). Dehydrated leaf tissue was transferred to a staining solution containing 1 % (w/v) Safranin-O (SigmaAldrich) in 100 % ethanol for 2 minutes. Leaf tissue was then washed in 100 % ethanol 3 times, and rehydrated, by repeating the dehydration series in the opposite order. Major veins were imaged using a flatbed scanner (CanoScan 9000F, Cannon LTD, UK) at 4800 × 4800 dpi resolution (Fig. 1). Minor veins were imaged using a camera (GXCAM HiChrome-S, GT Vision Ltd) attached to a light microscope (Leitz Diaplan). When measuring minor veins, images were acquired at three locations on the leaf tissue, as technical replicates. ImageJ (NIH) was used to measure VLA on images. Total leaf VLA was calculated as the sum of major and minor VLA. The total VLA was used to calculate intervein distance using the equation

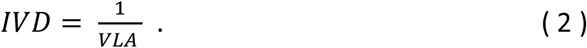

**Fig. 1:**
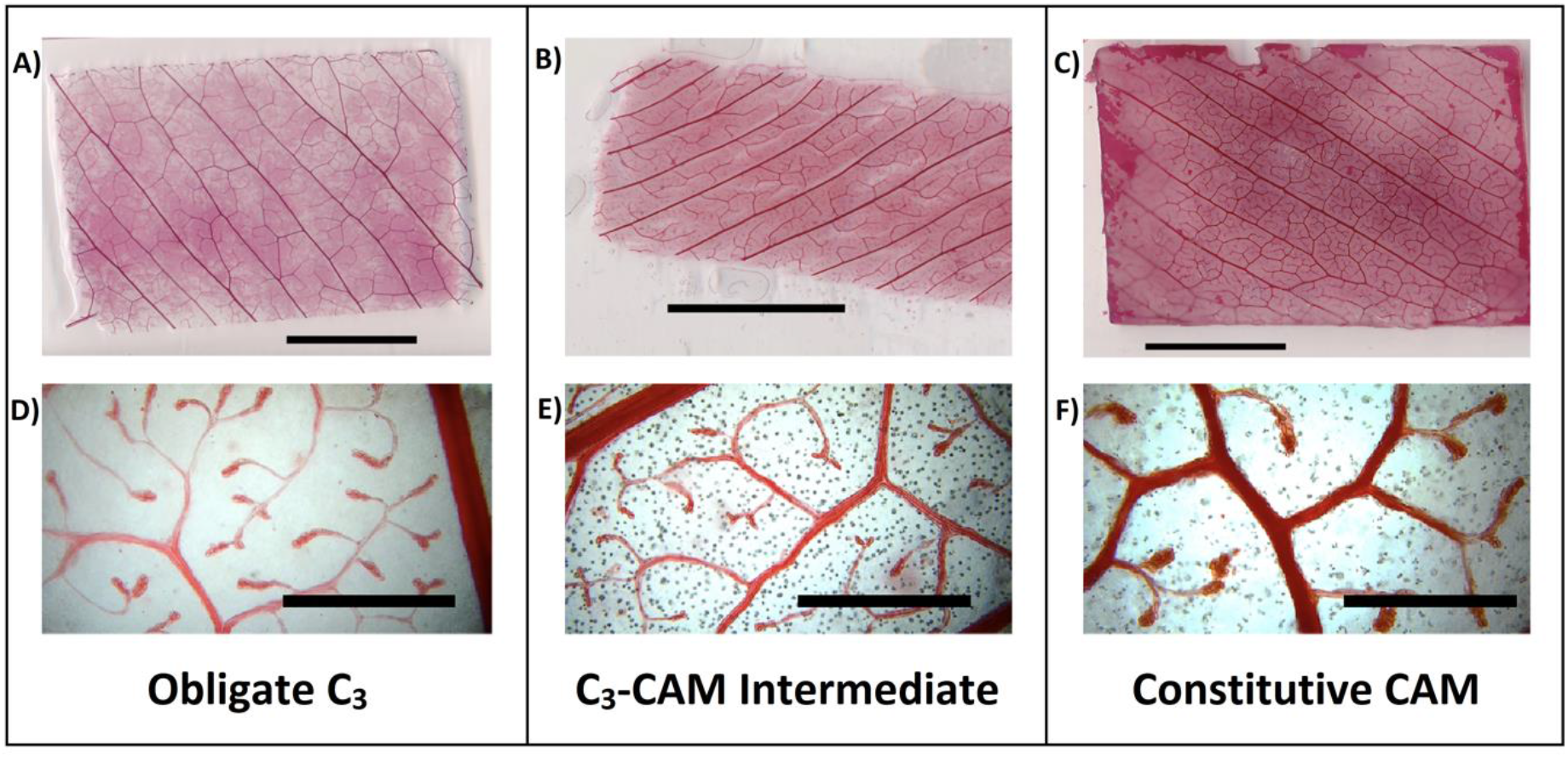
Example images used to estimate major and minor vein length per leaf area (VLA). Example of an obligate C_3_ species (*C. multiflora*) a C_3_-CAM intermediate (*C. pratensis*) and a constitutive CAM species (*C. alata*). (A-C) scale bar represents 10 mm. (D-F) Scale bar represents 1 mm.

The same leaf used to measure VLA was hand-sectioned to determine the average distance from vein to lower epidermal surface and the depth of hydrenchyma tissue. Hand-sectioned material was imaged using a camera (Q-IMAGING, QICAM, fast 1394) attached to a fluorescence microscope (Leica DMRB) under blue light. As veins are in one plane in *Clusia* leaves (Fig. S1), VED was calculated as the distance from the middle of a vein to the lower epidermal surface, averaged for 3 technical replicates, per leaf. For each species, 7-9 replicate leaves were used. *Clusia* leaves are hypostomatous, so the distance from veins to upper epidermis was not measured. Presence of a gel-like substance in *C. rosea* leaves prevented acquisition of clear images, so this species was omitted from this analysis (Fig. S2).

### Cross-Sectional Vessel Area

From each plant, the third or fourth leaf from the apex was sampled, and leaf area was measured using a flatbed scanner (hp Scanjet 5530 Photosmart Scanner, Hp, UK), as described above. Material used for sectioning was sampled from halfway along the proximal distal axis of the leaf, for midribs, as well as halfway along the petiole. Sectioning was done using a vibrating microtome (Ci LTD, UK) which was set to cut 100 μm sections at 80 Hz. Sections were transferred to 5 % phosphate buffered saline (Sigma-Aldrich) and stored overnight. Sections were imaged using a camera (Q-IMAGING, QICAM, fast 1394) attached to a florescence microscope (Leica DMRB) under blue light. Autofluorescence of lignin allowed visualisation of veins (Maceda and Terrazas, 2022). For each image, the cross-sectional vessel area was measured using ImageJ (NIH). For each species, n = 4, for both petiole and midribs.

### Statistics

All statistics were performed using R version 4.1.2.

## Results

### Hypothesis 1: Are CAM and/or elevated C_FT_ associated with lower VLA?

We analysed 10 species of *Clusia* to determine if CAM and/or elevated C_FT_ were associated with lower VLA. Interspecific comparisons found that species that do a greater proportion of their CO_2_ assimilation at night under well-watered conditions (CAM_ww_), had significantly lower total VLA (Fig. 2A). Separate analysis of the major and minor VLA found that only the latter significantly correlated with CAM_ww_ (Fig. 2C). No correlation was found between CAM_ww_ and major VLA (Fig. 2B). As minor VLA, and not major VLA, is lower in species that do CAM, the ratio of major to minor veins showed a significant positive correlation with CAM_ww_ (Fig. S3). In contrast, CAM_d_ did not correlate with total VLA (Fig. 2D) or minor VLA (Fig. 2F). However, CAM_d_ positively correlated with major VLA (Fig. 2E). Interspecific variation in C_FT_ did not correlate with total, major, or minor VLA (Fig. 2G-I). Taken together, these data suggest that low VLA is co-adapted with constitutive CAM physiology, not facultative CAM or elevated C_FT_. CAM_ww_ and C_FT_ are strongly associated with leaf and hydrenchyma depth, respectively (Barrera-Zambrano *et al*., 2014; Leverett *et al*., 2022). Consequently, leaf depth negatively correlated with total VLA and minor VLA (Fig. S4). In addition, no correlation was observed between hydrenchyma depth and total, major or minor VLA (Fig. S4).

**Fig. 2:**
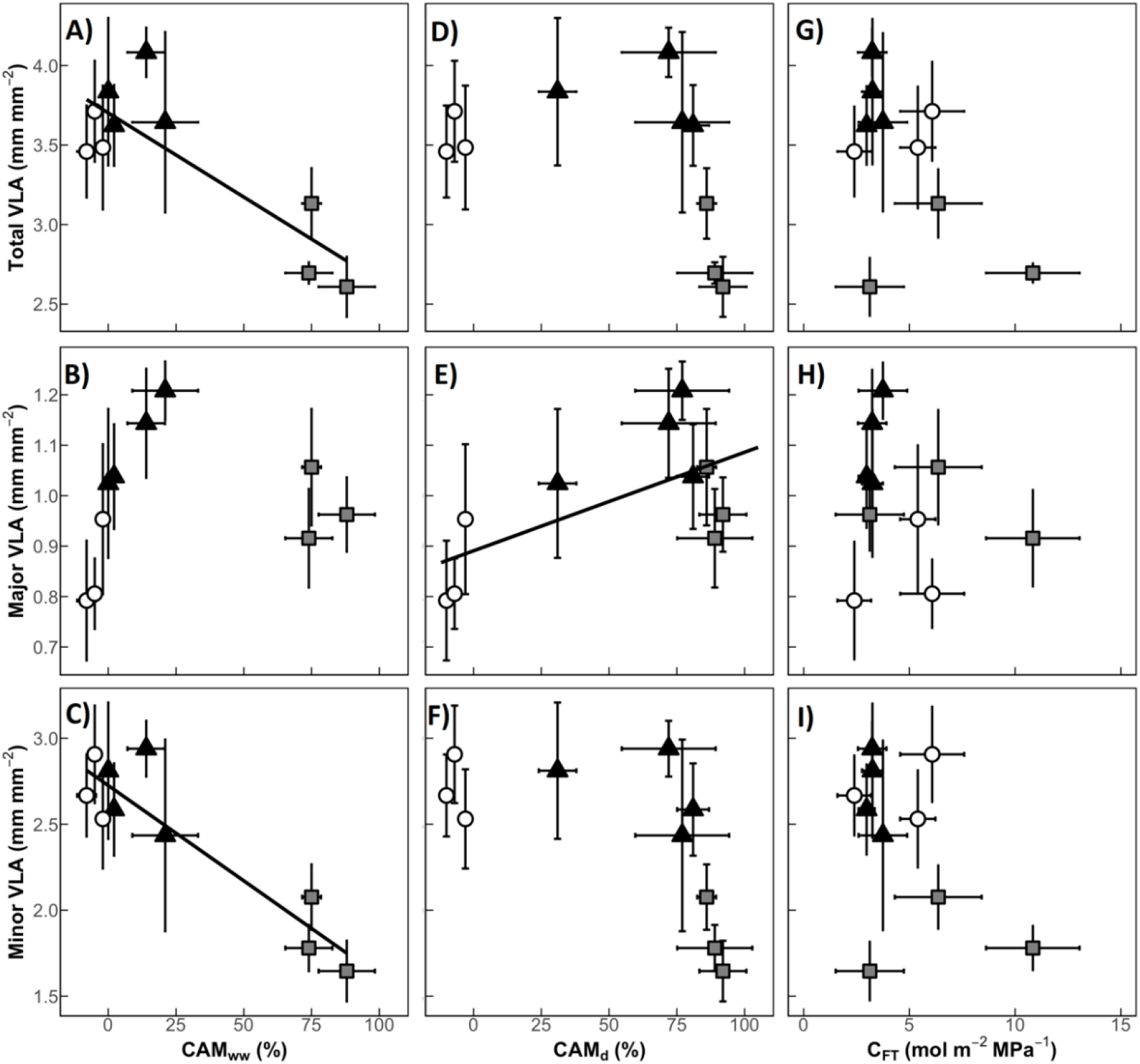
CAM species have lower vein length per leaf area (VLA), due to lower density of minor veins. **(A)** A negative correlation exists between the proportion of diel CO_2_ assimilation ocurring at night in well-watered plants (CAM_ww_) and total VLA (linear regression: R^2^ = 0.66, p = 0.003). (**B)** CAM_ww_ does not correlate with major VLA (linear regression: p = 0.69). **(C)** CAM_ww_ negatively correlates with minor VLA (linear regression: R^2^ = 0.80, p < 0.001). **(D)** The percentage of CO_2_ assimilation occurring at night in drought-treated plants (CAM_d_) does not correlate with total VLA (linear regression: p = 0.24). **(E)** CAM_d_ positively correlates with major VLA (linear regression: R^2^ = 0.33, p = 0.05). **(F**) CAM_d_ does not correlate with minor VLA (p = 0.06). **(G)** Bulk hydraulic capacitance (C_FT_) does not correlate with total VLA (linear regression: 0.15). **(H)** C_FT_ does not correlate with major VLA (linear regression: 0.53). **(I)** C_FT_ does not correlate with minor VLA (linear regression: 0.19). White circles = obligate C_3_ species, black triangles = C_3_-CAM intermediates, grey squares = constitutive CAM species. Error bars are ± 1 standard deviation and for each species, n = 7-9.

Minor veins often end at vein termini (Fig. 1D-F), at which point all water moves out of the xylem and into the mesophyll tissue. We suspected that minor VLA and vein termini density (VTD) would positively correlate with each other. Analysis of 10 *Clusia* species confirmed this hypothesis, as minor VLA correlated significantly with VTD (Fig. 3A). In addition, CAM_ww_ negatively correlated with VTD (Fig. 3B). No correlation was found between C_FT_ and VTD. Furthermore, leaf depth negatively correlated with VTD (Fig. S5), whereas hydrenchyma depth and VTD did not correlate (Fig. S5).

**Fig. 3:**
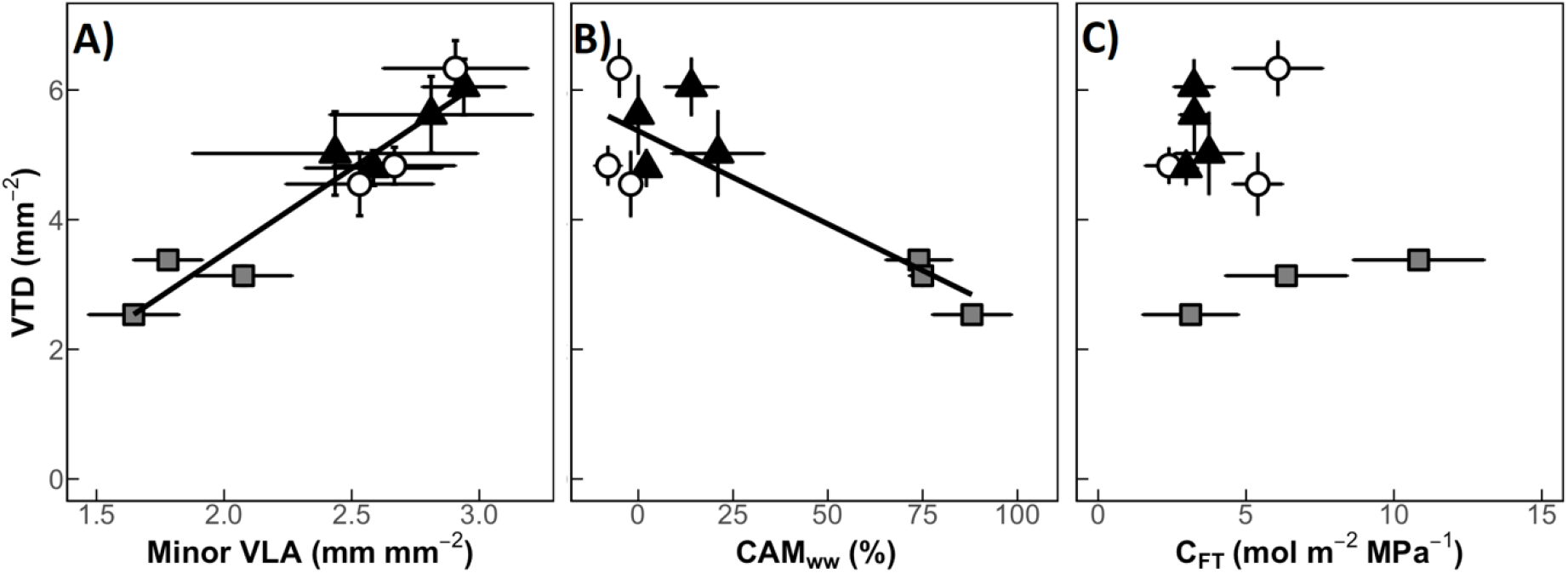
CAM species have lower leaf vein termini density (VTD), due to their lower minor vein length per leaf area (minor VLA). **(A)** Minor VLA positively correlates with VTD (linear regression: R^2^ = 0.90, p < 0.001). **(B)** The proportion of diel CO_2_ assimilation ocurring at night (CAM_ww_) negatively correlates with VTD (linear regression: R^2^ = 0.71, p = 0.001) **(C)** Bulk hydraulic capacitance (C_FT_) does not correlate with VTD (linear regression: p = 0.37). White circles = obligate C_3_ species, black triangles = C_3_-CAM intermediates, grey squares = constitutive CAM species. Error bars are ± 1 standard deviation and for each species, n = 7-9.

### Hypothesis 2: Are CAM and/or elevated C_FT_ associated with IVD:VED ratios < 1?

Unlike the majority of angiosperms, some *Clusia* species have IVD:VED ratios < 1: a phenomenon termed ‘vascular overinvestment’ (Fig. 4). Across 10 species of *Clusia* IVD:VED ratios positively correlated with CAM_ww_. In contrast no correlation was observed between C_FT_ and IVD:VED ratios. In addition, IVD:VED ratios positively correlated with leaf depth, but did not correlate with hydrenchyma depth (Fig. S6).

**Fig. 4:**
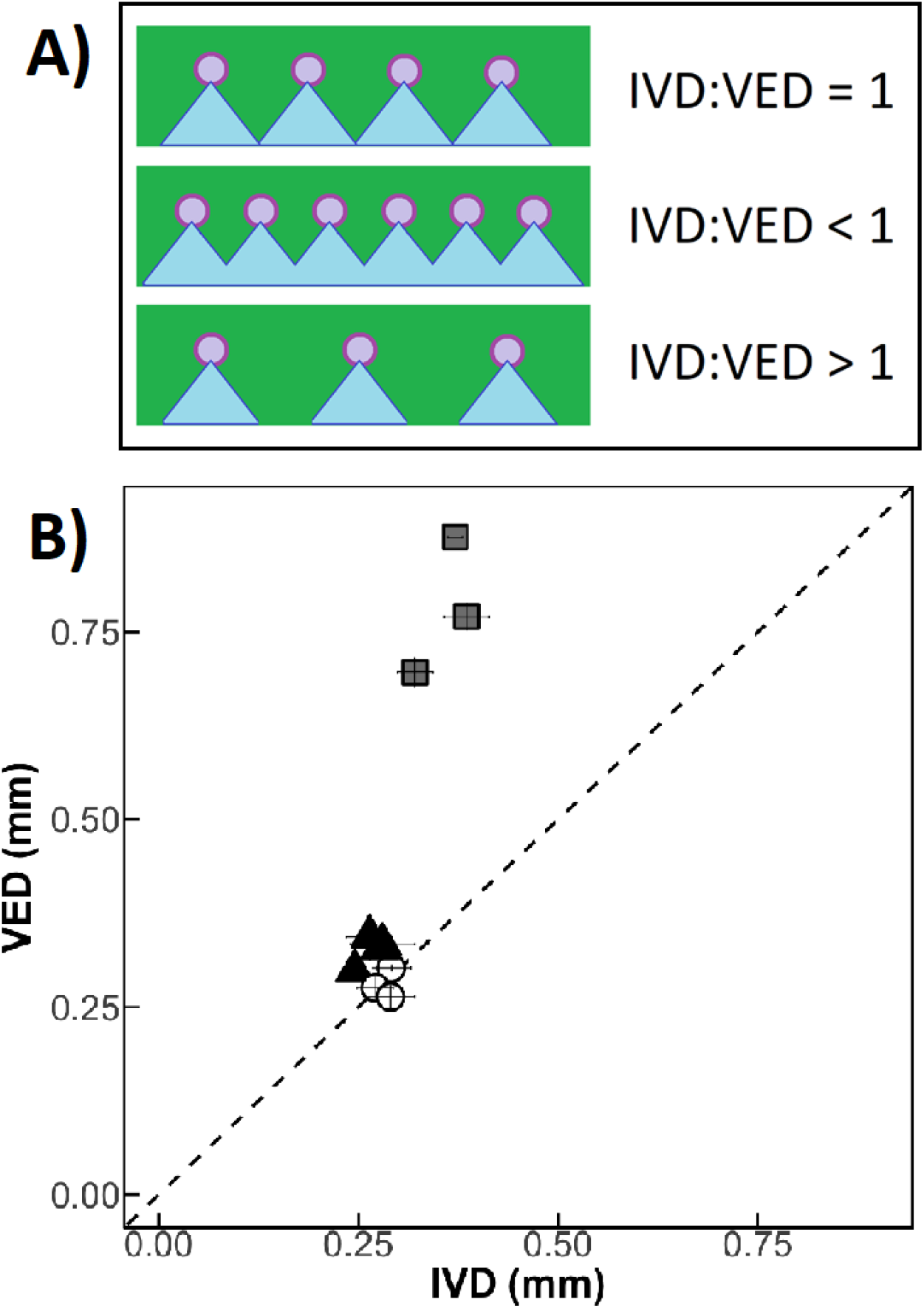
Coordination of intervein distance (IVD) and vein to lower epidermal distance (VED) in leaves. **(A)** In most angiosperm species, optimal placement of veins (purple), in leaves (green) is one where IVD:VED = 1. In this scenario, water (blue) can permeate efficiently into the abaxial mesophyll tissue. When IVD:VED < 1, superfluous veins are present, which do not increase the efficiency with which water permeates into the abaxial mesophyll. When IVD:VED > 1, insufficient veins are present to replace efficiently replace water lost via abaxial stomata. **(B)** Across 10 species of *Clusia*, species exist where IVD:VED < 1. White circles = obligate C_3_ species, black triangles = C_3_-CAM intermediates, grey squares = constitutive CAM species. Error bars are ± 1 standard deviation and for each species, n = 7-9.

### Hypothesis 3: Are CAM and/or elevated C_FT_ associated with wider vessels in the midrib and petiole?

The diameter of vessels is known to impact vulnerability to embolism when Ψ_L_ becomes more negative: as larger vessels more readily seed bubbles that break the transpiration stream. Since CAM and elevated C_FT_ both slow the rate at which Ψ_L_ falls during drought events, we hypothesised that species with these adaptations may develop wider vessels, to increase efficient flow of water into leaves. However, across 11 species of *Clusia* neither CAM_ww_, CAM_d_ nor CFT correlated with cross-sectional vessel area in the midrib or petiole (Fig. 7). Cross-sectional vessel area in the midrib and petiole both correlated with leaf area (Fig. 7D,H). Taken together, these data suggest that the development of wider vessels is coordinated with leaf size, but not with CAM or C_FT_.

## Discussion

### Leaf Vascular Architecture Is Coordinated With Constitutive Crassulacean Acid Metbolism

Both CAM and elevated C_FT_ reduce the rate at which water moves through leaves. However, in *Clusia*, VLA appears to have been optimised to match the reduced hydraulic demands conferred by CAM, rather than elevated C_FT_. *Clusia* species with constitutive CAM will experience relatively low transpiration rates over the entire year, during both the dry and wet seasons (Holtum *et al*., 2004; Leverett *et al*., 2021). By sustaining low transpirational rates, constitutive CAM species require less water to move through the xylem in order to keep leaves hydrated. As constitutive CAM species require lower hydraulic conductance, they develop lower VLA and VTD than C_3_ relatives, thereby reallocating space and resources for the development of light-harvesting photosynthetic mesophyll cells. Low VLAs and VTDs are achieved by developing lower minor VLA (Fig. 2C), as minor veins are less structurally reinforced and more vulnerable to physical damage during drought (Sack and Scoffoni, 2013). In contrast, whilst the C_3_-CAM *Clusia* species can facultatively switch on or upregulate CAM during the dry season, they spend most of the year predominantly doing C_3_ photosynthesis, which will result in higher transpiration rates. Interestingly, C_3_-CAM species did not have intermediate values of total VLA or VTD, but instead their total VLA and VTD values more closely resembled those of obligate C_3_ species (Fig. 2A). It seems that C_3_-CAM species develop higher VLAs and VTDs in order to tolerate the greater rates of transpiration that they will experience most of the year, rather than to optimise vascular architecture for acute dry seasons when they switch to CAM. However, C_3_-CAM species did exhibit higher major VLA (Fig. 2E). Structurally reinforced major veins are thought to be less vulnerable to implosion and physical damage during drought. It is possible that higher major VLAs may serve as a solution for facultative CAM species: high minor VLA is optimised for transpiration rates associated with C_3_ physiology, alongside higher major VLA which helps to cope with low water potentials during hotter dry seasons.

In contrast to CAM, C_FT_ did not correlate with either VLA or VTD (Fig. 2–3). It is likely that the presence of CAM in the genus *Clusia* largely mitigates the need for leaves to co-adapt their VLA for elevated C_FT_. Put simply, CAM appears to have a greater influence than elevated C_FT_ on the architecture of xylem in leaves. Consequently, despite *Clusia* leaves exhibiting > 5-fold variation in C_FT_, this has very little influence on VLA or VTD. This finding is congruent with recent work comparing the effects of CAM and elevated C_FT_ on plant physiology during drought (Leverett *et al*., 2021, 2022). Ecophysiological modelling has demonstrated that, in *Clusia*, the presence of CAM largely obviates the effect of elevated C_FT_ on transpiration rates (Leverett *et al*., 2022). As CAM dampens the effect of elevated C_FT_ on leaf hydraulic physiology the former is likely to have a more substantive influence on xylem anatomy. In addition, from an ecological perspective, CAM determines the distribution of *Clusia* species across a precipitation gradient, whereas hydrenchyma depth (which confers elevated C_FT_) does not (Leverett *et al*., 2021). The data presented in this study reinforce these ecological patterns: it appears that CAM, and not C_FT_ is the key hydraulic innovation that influences the ecology, physiology and leaf vascular anatomy of *Clusia*.

We also estimated IVD:VED ratios across species of *Clusia* to understand how vein density is coordinated with leaf depth. In contrast to most angiosperms (Zwieniecki and Boyce, 2014), some species of *Clusia* had IVD:VED ratios < 1 (Fig. 4). Similar ‘vascular over investment’ has been observed in other taxa and appears to be associated with thicker leaves. However, it has been unclear if vascular over investment is an adaptation to overcome the longer apoplastic distance imposed by greater leaf depth, or if IVD:VED ratios < 1 occur in order to facilitate quick refilling of water stores that provide C_FT_ (de Boer *et al*., 2016; Males, 2017a). In *Clusia*, CAM/leaf depth are independent of C_FT_/hydrenchyma depth, making this genus ideal for exploring these opposing hypotheses (Barrera-Zambrano *et al*., 2014; Leverett *et al*., 2022). Our data show that IVD:VED ratios correlate with CAM and leaf depth (Fig. 5, S6), and not with C_FT_ or hydrenchyma depth. In CAM species, thick, succulent photosynthetic chlorenchyma tissue is required to provide adequate space for nocturnal storage of malic acid (Barrera-Zambrano *et al*., 2014; Males, 2018; Töpfer *et al*., 2020). However, this will increase the apoplastic distance through which water must travel, in order to infiltrate the entire leaf. As a result, it is likely that CAM species require IVD:VED ratios < 1, in order to provide this additional water to the mesophyll and keep the leaf hydrated when stomata are open.

**Fig. 5:**
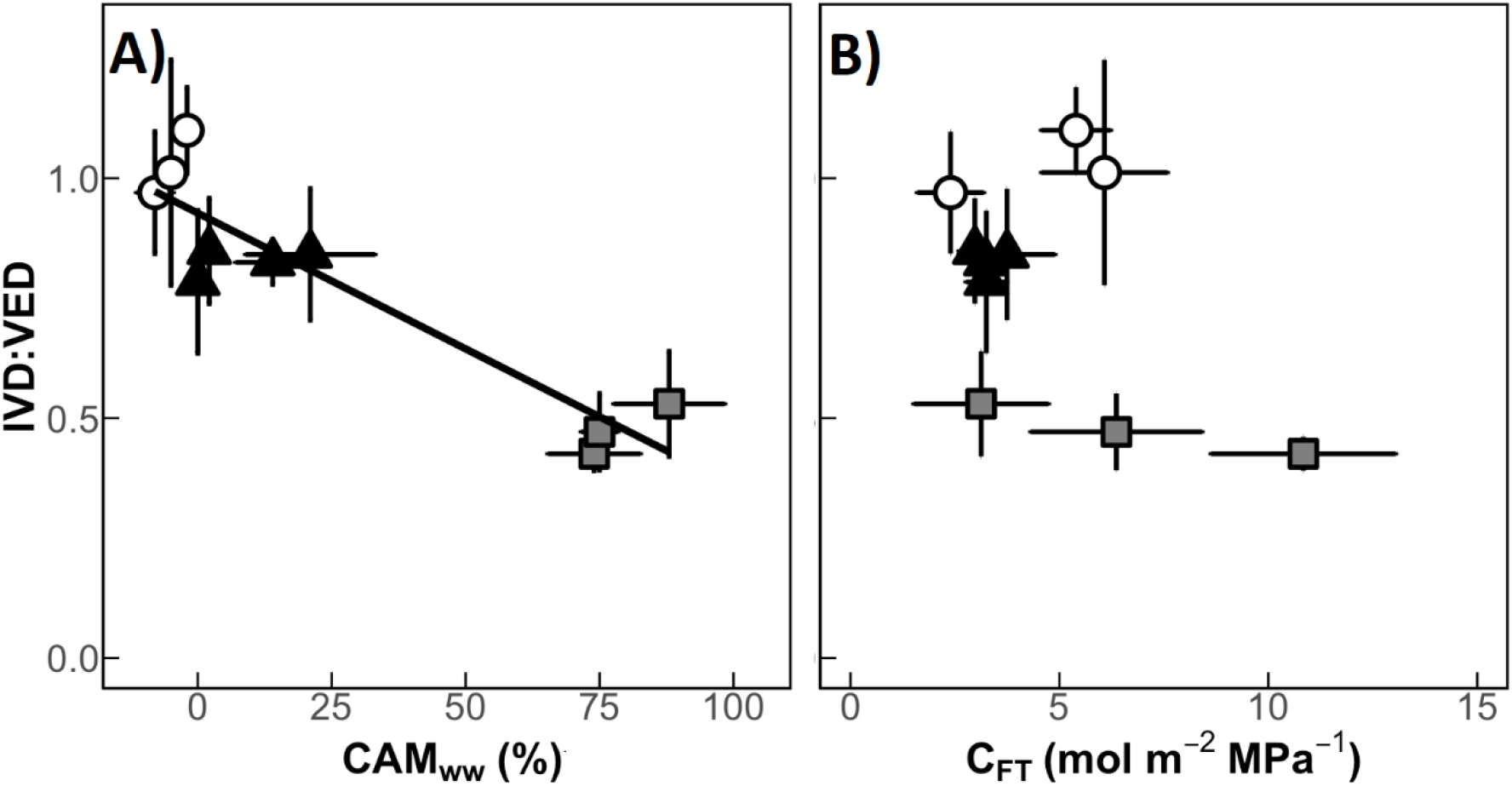
The ratio of intervein distance and vein to epidermis distance (IVD:VED) is higher in CAM species than C_3_ species. **(A)** The proportion of diel CO_2_ assimilation ocurring at night (CAM_ww_) negatively correlates with IVD:VED ratios across 10 species of *Clusia* (linear regression: R^2^ = 0.83, p < 0.001). **(B)** Bulk hydraulic capacitance (C_FT_) does not correlate with IVD:VED ratios (linear regression: p = 0.18). White circles = obligate C_3_ species, black triangles = C_3_-CAM intermediates, grey squares = constitutive CAM species. Error bars are ± 1 standard deviation and for each species, n = 7-9.

**Fig. 6:**
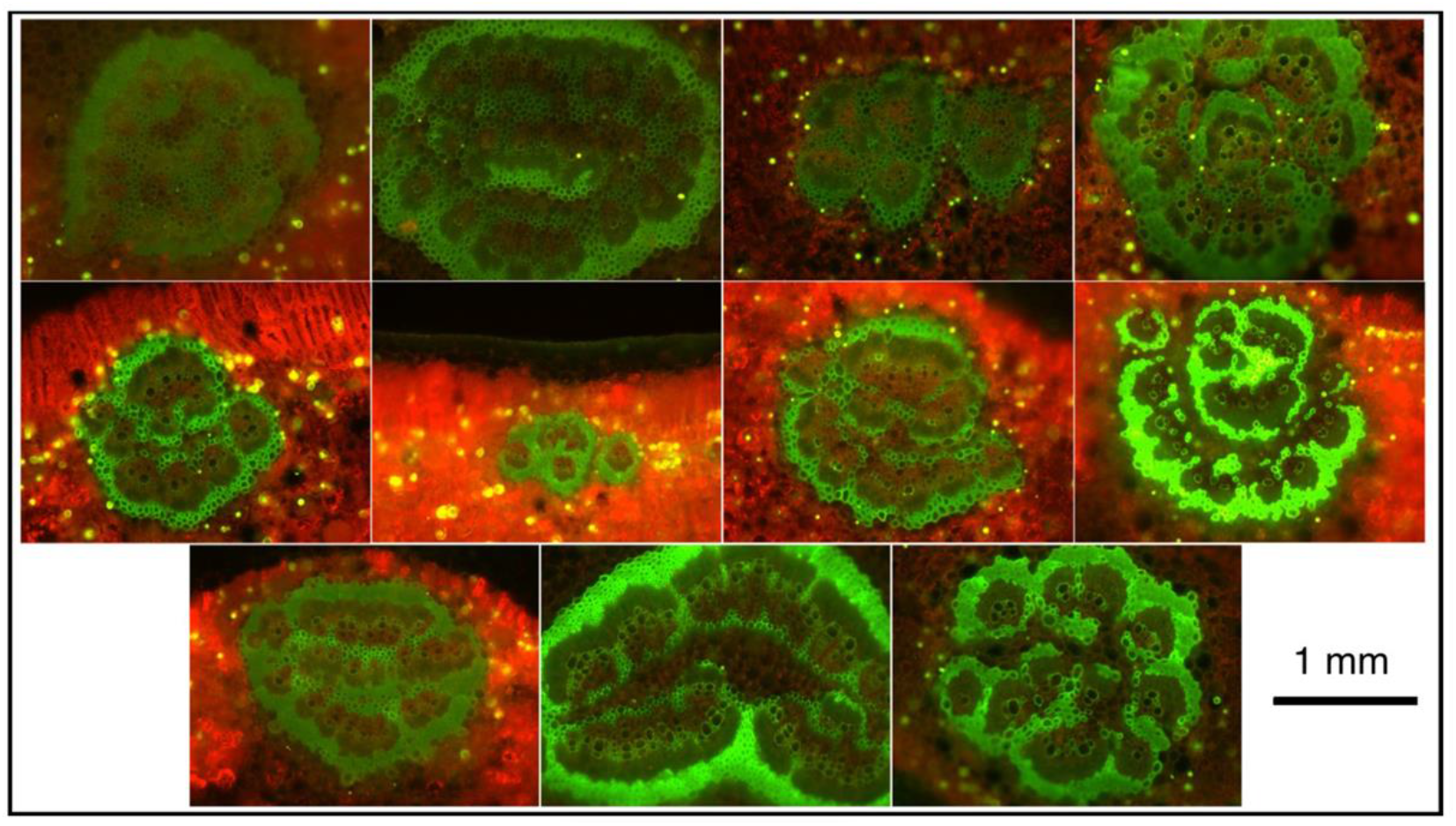
Example images used to measure midrib xylem vessels. Top row (constitutive CAM species), from left to right: *Clusia hilariana, C. fluminensis, C. alata, C. rosea*. Middle row (C_3_-CAM intermediates): *C. minor, C. lanceolata, C. pratensis, C. aripioensis*. Bottom row (obligate C_3_ species): *C. tocuchensis, C. multiflora, C. grandiflora*.

**Fig. 7:**
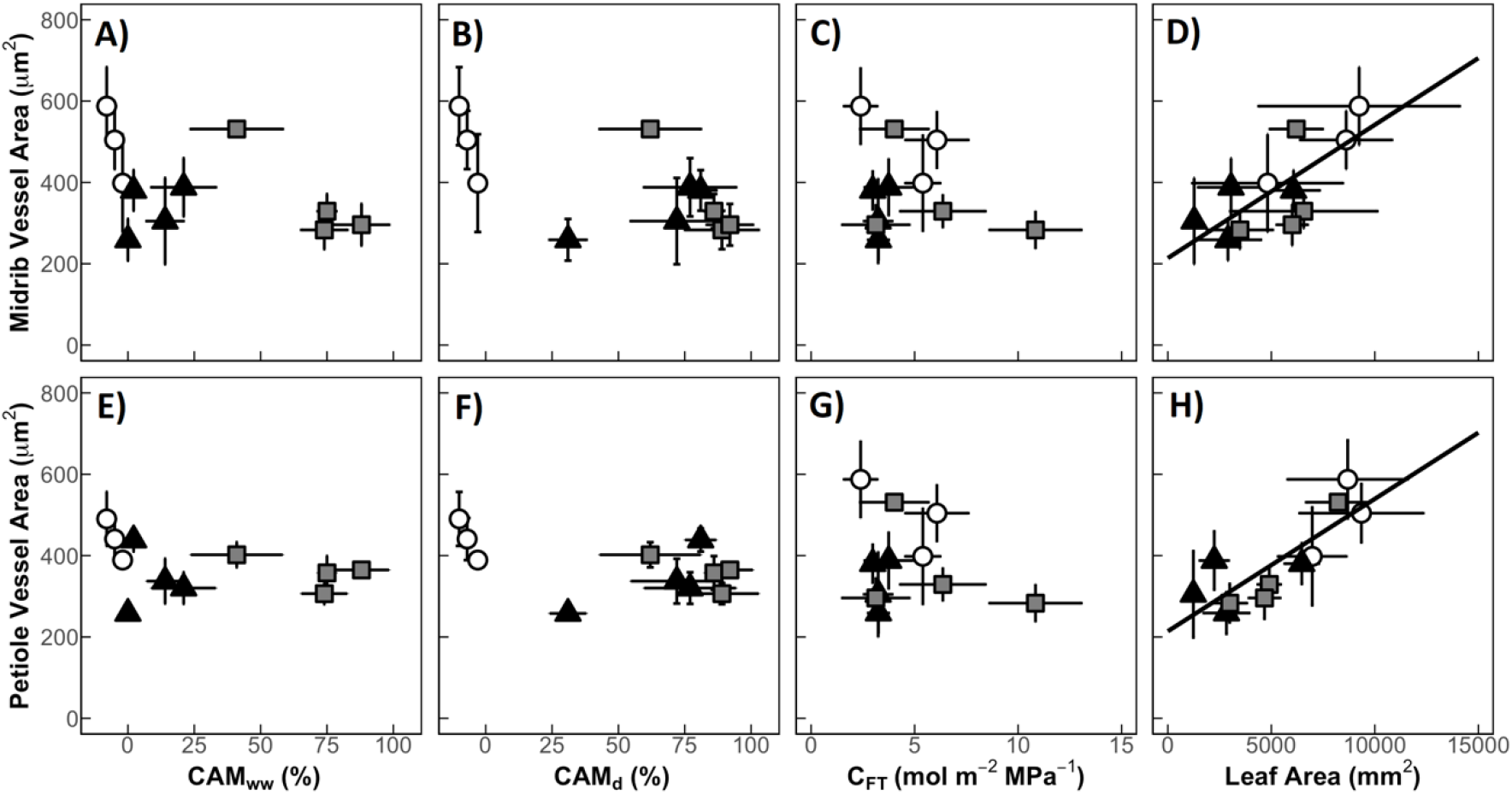
Neither CAM nor C_FT_ correlate with vessel cross-sectional area. **(A)** The proportion of diel CO_2_ assimilation occurring at night in well-watered plants (CAM_ww_) does not correlate with midrib cross-sectional vessel area (linear regression: p = 0.17). **(B)** The proportion of diel CO_2_ assimilation ocurring at night in drought-treated plants (CAM_d_) does not correlate with midrib cross-sectional vessel area (linear regression: p = 0.051). **(C)** Bulk hydraulic capacitance (C_FT_) does not correlate with midrib cross-sectional vessel area (linear regression: p = 0.46). **(D)** Leaf area positively correlates with midrib cross-sectional vessel area (linear regression: R^2^ = 0.49, p = 0.01). **(E)** CAM_ww_ does not correlate with petiole cross-sectional vessel area (linear regression: p = 0.30). **(F)** CAM_d_ does not correlate with petiole cross-sectional vessel area (linear regression: p = 0.15). **(G)** C_FT_ does not correlate with midrib cross-sectional vessel area (linear regression: p = 0.42). **(H)** Leaf area positively correlates with midrib cross-sectional vessel area (linear regression: R^2^ = 0.70, p < 0.001). White circles = obligate C_3_ species, black triangles = C_3_-CAM intermediates, grey squares = constitutive CAM species. Error bars are ± 1 standard deviation and for each species, n = 4.

### Learning From Nature: Xylem Traits Should Be Optimised For CAM Biodesign

Due to the effects of global warming, aridity will increase across much of the worlds arable land. Therefore, it is integral that scientists develop novel approaches to increase drought tolerance in crops (Borland *et al*., 2015; Cushman *et al*., 2015; Leakey *et al*., 2019, Pan *et al*., 2021). To this end, CAM plants have a great deal of potential, as their low transpiration rates and elevated water use efficiency allow them to tolerate conditions in marginal land that is less amenable to C_3_ or C_4_ crops species (Borland *et al*., 2014). Furthermore, as the majority of crops do not do CAM, considerable efforts are underway to bioengineer this pathway into C_3_ species, to prepare for hotter, drier futures. Beyond introducing the key enzymes that comprise the CAM pathway (Lim *et al*., 2019), it is essential that the anatomy of host plants is appropriate for CAM to function efficiently. To this end, the introduction of a grape helix-loop-helix transcription factor (*VvCEB1*) has been shown to increase cell size in *Arabidopsis thaliana*, which will provide adequate space for nocturnal accumulation of malate (Lim *et al*., 2018, 2020). However, little attention has been given to the xylem adaptations associated with CAM, and how these vascular traits might be optimised to maximise water use efficiency in bioengineered CAM plants (Borland *et al*., 2018). The data presented in this study suggest that low VLA and VTD, alongside IVD:VED ratios < 1 should be selected when choosing a host plant for CAM bioengineering. It is possible that the *35S::VvCEB1* overexpression lines engineered by Lim *et al*., (2018) exhibit lower VLA and/or higher IVD:VED ratios, due to their wider leaves and larger mesophyll cells. However, xylem anatomy remains unreported for these transgenic plants. If the optimal vascular architecture is not found in *35S::VvCEB1* plants, other manipulations could be incorporated to achieve this goal, such as using leaf-specific promoters to manipulate the auxin signalling pathway (Perico *et al*., 2022). For example, upregulating cyclophilin 1 cis/trans isomerase (CYP1), alongside a *35S::VvCEB1*, could be used to remove repression of auxin response factors (ARFs) during leaf development, and generate lower VLAs (Andrade *et al*., 2022). Future bioengineering efforts must look beyond mesophyll anatomy, to ensure optimal vascular traits are introduced alongside the core enzymes that catalyse the CAM cycle.

### *Conclusions – Towards a Complete Anatomical Characterisation of* Clusia

The data presented here represent the first characterisation of vascular anatomy associated with CAM photosynthesis at the taxonomic level of the genus. The present study demonstrates that CAM is not only associated with the anatomy of photosynthetically active tissues in *Clusia* (Barrera-Zambrano *et al*., 2014; Luján, *et al*., 2021), but also coordinated with vascular architecture; the low transpiration rates resulting from CAM are associated with low VLAs, and the succulent photosynthetic tissue appears to require an high IVD:VED ratios to efficiently provide water to the mesophyll. These findings indicate that CAM requires a complex suite of anatomical changes, to optimise both the photosynthetic and hydraulic needs of the leaf.

## Acknowledgements

This research was funded by Newcastle University’s R. B. Cook Scholarship, a Smithsonian Tropical Research Institute Fellowship grant to AL and a Newcastle University Vacation Scholarship to KF. We would also like to thank Samuel Logan and Helen Martin for their assistance with *Clusia* propagation/cultivation and transportation of material from Cockle Park to Newcastle Campus.

## Supplementary Figures

**Fig S1:**
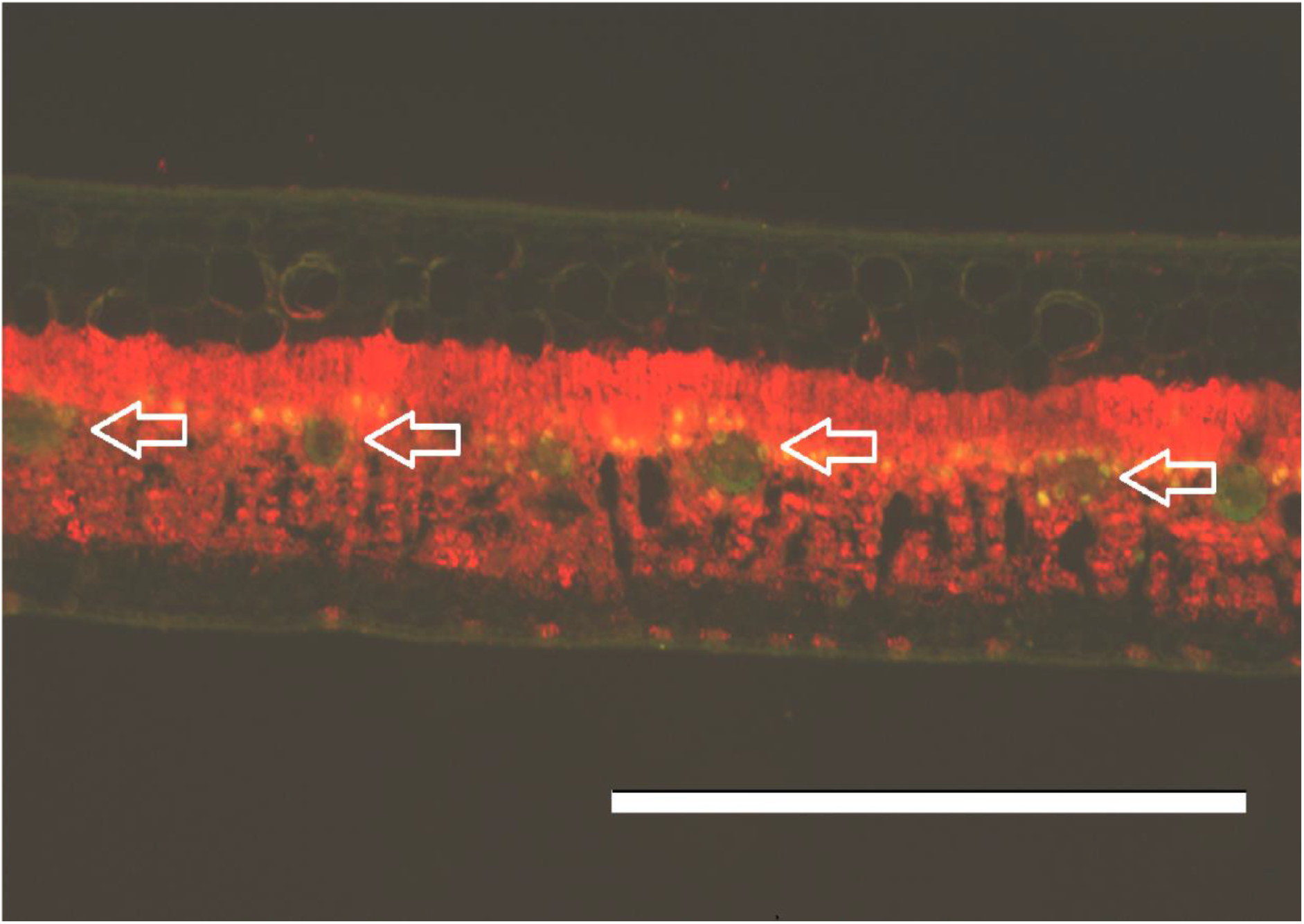
Veins develop in one plane in *Clusia* leaves. Image shows the mesophyll of *C. tocuchensis*, with arrows pointing to the veins, which all fall in one flat plane in the leaf. All species in this study had veins in one, flat plane between the palisade and spongy mesophyll tissue layers. Scale bar = 1 mm.

**Fig. S2:**
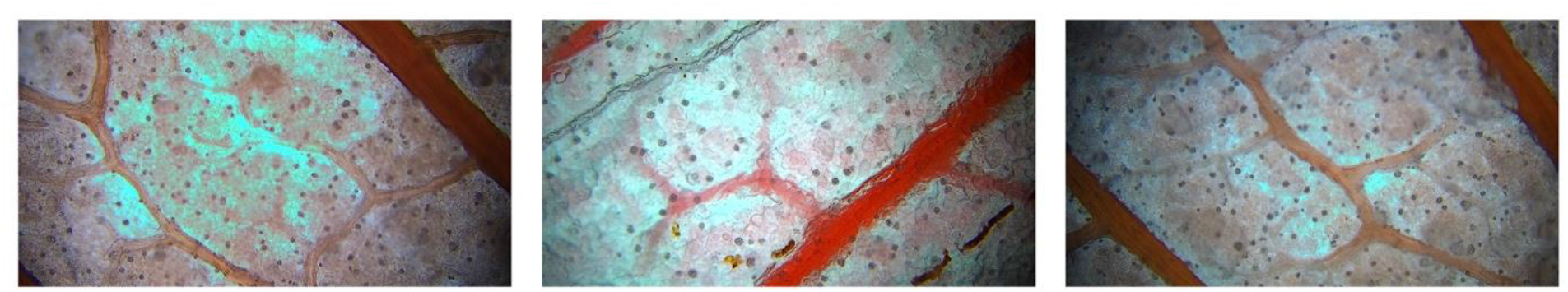
Gel-like substance prevented clear vein density images from being acquired for *Clusia rosea*. Higher order veins were visible but smaller, lower order veins were obscured.

**Fig. S3:**
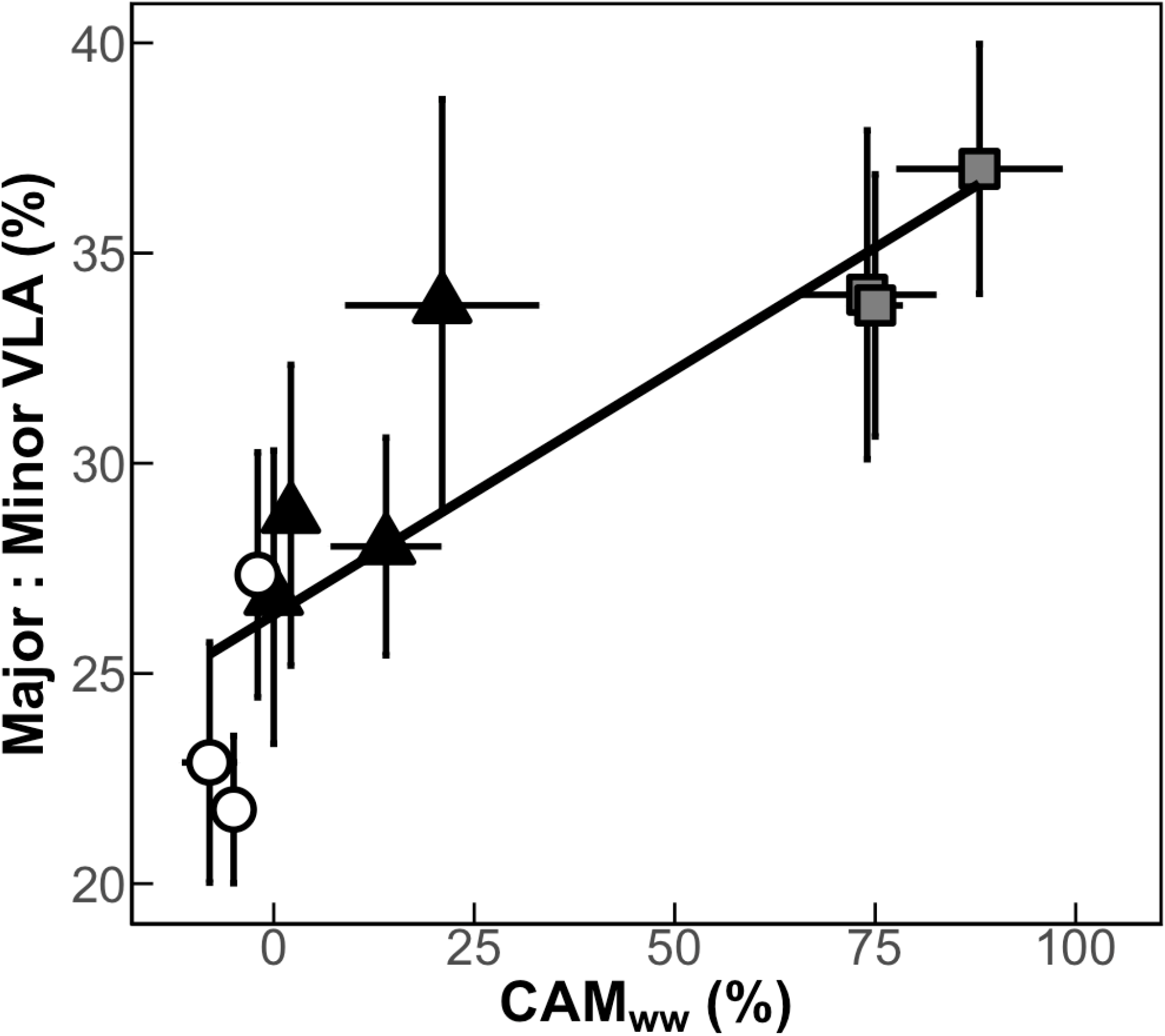
Major veins make up a greater proportion of total VLA in CAM species. The proportion of diel CO_2_ assimilation occurring at night in well-watered plants (CAM_ww_) positively correlates with the percentage of total VLA comprised of major veins (Major : Minor VLA) (linear regression: R^2^ = 0.73, p = 0.001). White circles = obligate C_3_ species, black triangles = C_3_-CAM intermediates, grey squares = constitutive CAM species. Error bars are ± 1 standard deviation and for each species, n = 7-9

**Fig. S4:**
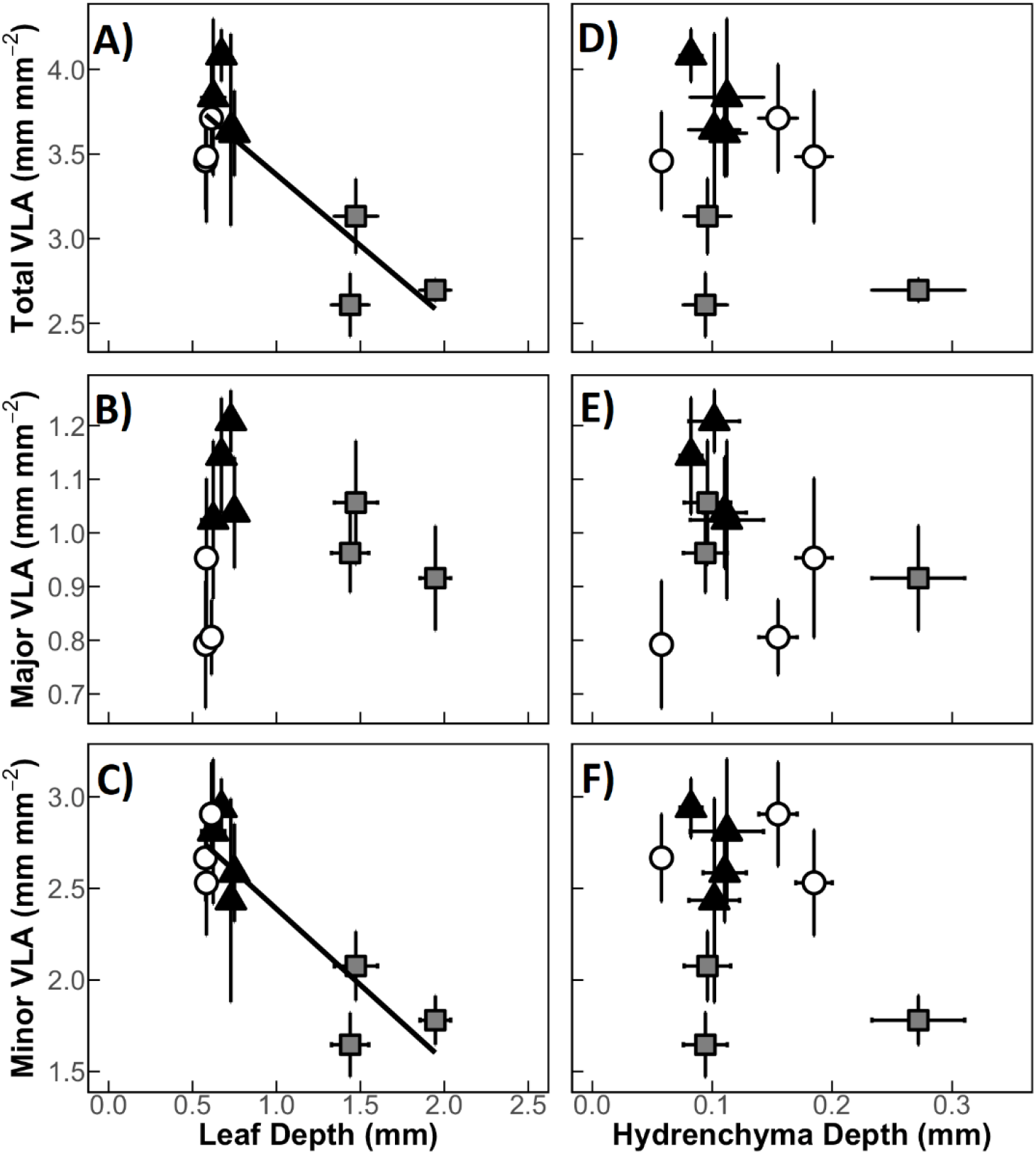
Relationships between VLA, leaf depth and hydrenchyma depth across 10 *Clusia* species. **(A)** A negative correlation exists between the leaf depth and total VLA (linear regression: R^2^ = 0.71, p = 0.001). (**B)** Leaf depth does not correlate with major VLA (linear regression: p = 0.94). **(C)** Leaf depth negatively correlates with minor VLA (linear regression: R^2^ = 0.77, p < 0.001). **(D)** Hydrenchyma depth does not correlate with total VLA (linear regression: 0.26). **(E)** Hydrenchyma depth does not correlate with major VLA (linear regression: 0.48). **(F)** Hydrenchyma depth does not correlate with minor VLA (linear regression: 0.35). White circles = obligate C_3_ species, black triangles = C_3_-CAM intermediates, grey squares = constitutive CAM species. Error bars are ± 1 standard deviation and for each species, n = 7-9

**Fig. S5:**
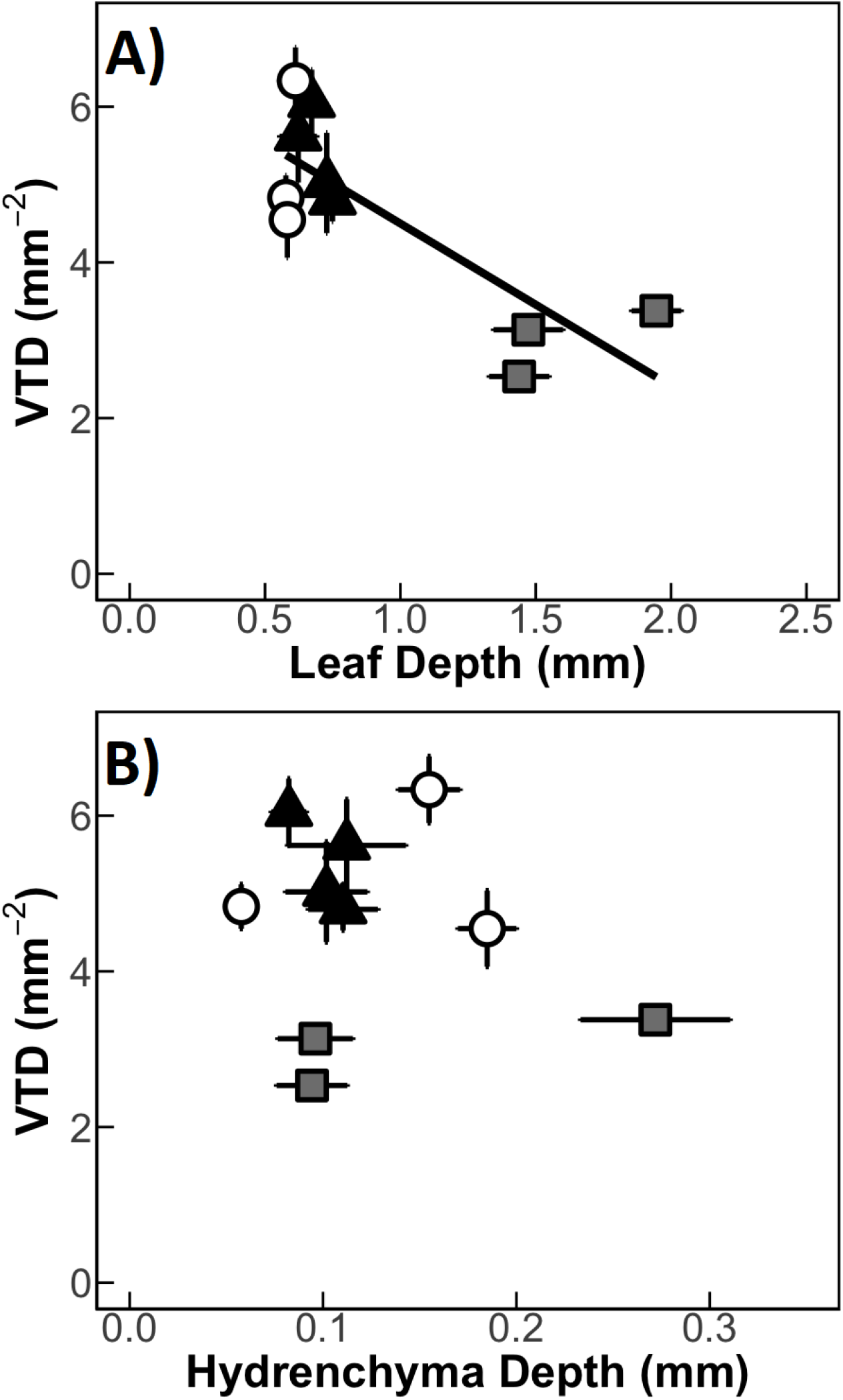
Relationship between leaf vein termini density (VTD), leaf depth and hydrenchyma depth across 10 species of *Clusia*. **(A)** Leaf depth negatively correlates with VTD (linear regression: R^2^ = 0.61, p = 0.004) **(B)** Hydrenchyma depth does not correlate with VTD (linear regression: p = 0.62). White circles = obligate C_3_ species, black triangles = C_3_-CAM intermediates, grey squares = constitutive CAM species. Error bars are ± 1 standard deviation and for each species, n = 7-9.

**Fig. S6:**
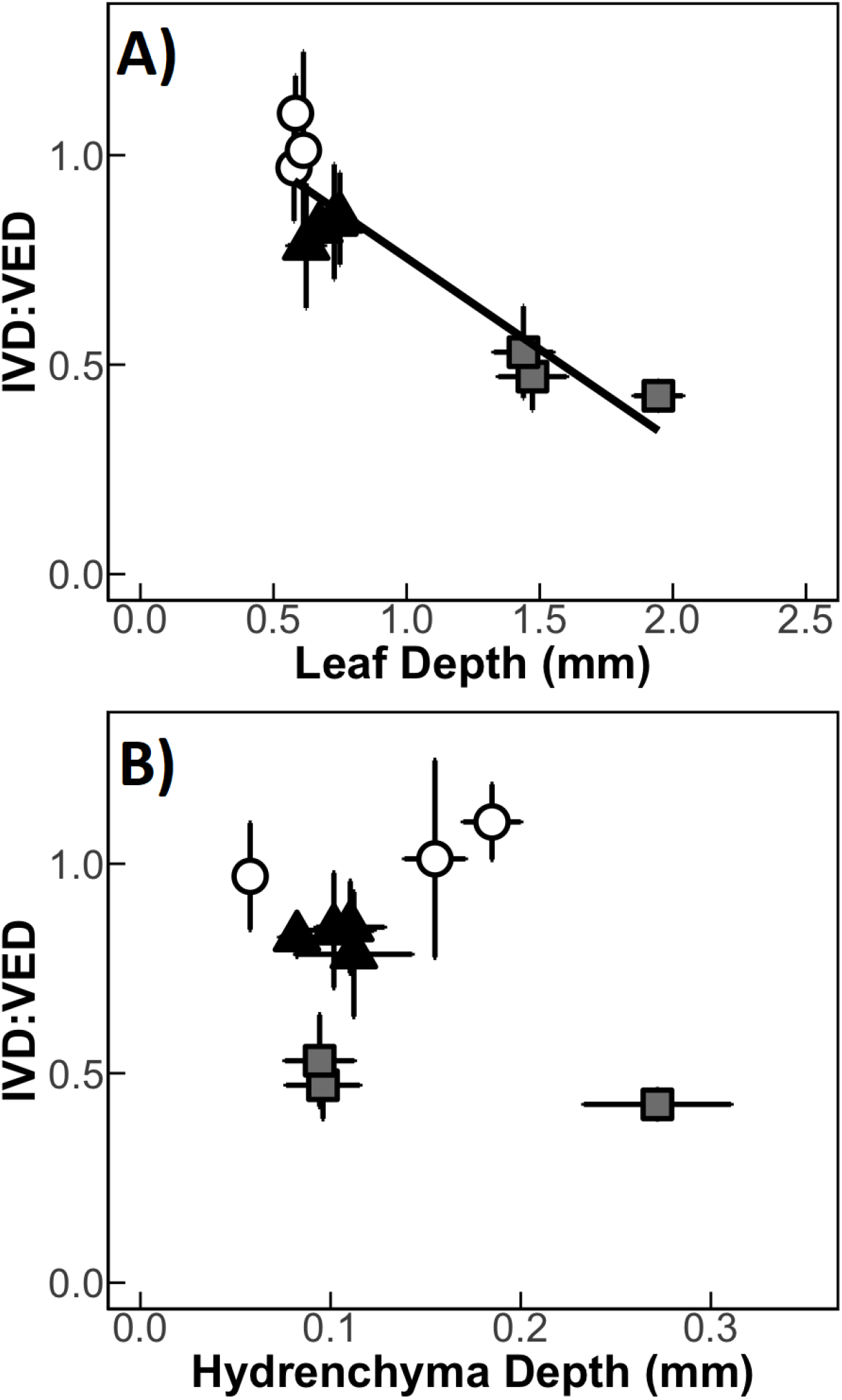
Relationship between IVD:VED ratios, leaf depth and hydrenchyma depth across 10 species of *Clusia*. **(A)** Leaf depth negatively correlates with IVD:VED ratio (linear regression: R^2^ = 0.83, p < 0.001). **(B)** Hydrenchyma depth does not correlate with IVD:VED ratio (linear regression: p = 0.57). White circles = obligate C_3_ species, black triangles = C_3_-CAM intermediates, grey squares = constitutive CAM species. Error bars are ± 1 standard deviation and for each species, n = 7-9.

